# Augmented efficacy of uttroside B over sorafenib in a murine model of human hepatocellular carcinoma

**DOI:** 10.1101/2022.02.01.478749

**Authors:** Mundanattu Swetha, C.K. Keerthana, Tennyson P Rayginia, Lekshmi R Nath, Nair Hariprasad Haritha, Kalishwaralal Kalimuthu, Somaraj Janet, Sreekumar Pillai, Kuzhuvelil B Harikumar, Sankar Sundaram, Nikhil Ponnoor Anto, Dee H Wu, Ravi Shankar Lankalapalli, Rheal Towner, Noah Isakov, Deepa Sathyaseelan, Ruby John Anto

## Abstract

**Background:** We previously reported the potency of *S. nigrum*-derived uttroside B (Utt-B). Recently Utt-B is flagged as an ‘orphan drug’ against hepatocellular carcinoma (HCC) by the US FDA. The current study aims to validate the enhanced *in vivo* efficacy of Utt-B over sorafenib, the first-line treatment option against HCC.

**Methods:** Human liver cancer cell line, HepG2 was employed as an HCC model and the comparison between Utt-B *vs* sorafenib therapeutic efficacies against HCC *in vivo* were evaluated in NOD.CB17-Prkdcscid/J mice that bear HepG2-induced HCC xenografts.

**Results:** Our data indicate that Utt-B is a more potent anti-HCC drug than sorafenib, *in vivo*. Apart from the superior therapeutic benefit over sorafenib, Utt-B is pharmacologically safer *in vivo*, and owing to this virtue, the drug-induced side effects are largely alleviated in the context of HCC chemotherapy.

**Conclusions:** Our data demonstrate the superior therapeutic index of Utt-B over sorafenib against HCC. Clinical studies in HCC patients utilizing Utt-B, which now holds the US FDA approval as an ‘orphan drug’, is an essential step to promote this drug from bench to bedside.

---

According to the GLOBOCAN database, liver cancer is the third leading cause of cancer death worldwide in 2020, with approximately 830,000 deaths (Sung, Ferlay et al. 2021). Sorafenib, the FDA-approved drug for HCC, is an orally administered multi-kinase inhibitor that inhibits cell proliferation via the inhibition of the serine/threonine kinase RAF (Llovet, Ricci et al. 2008). Moreover, sorafenib targets pro-angiogenic VEGF and PDGFR (Ibrahim, Yu et al. 2012). Sorafenib is widely studied and numerous data, including the clinical trials, reveal the legitimate reason for this drug to be administered as a first-line option to treat HCC patients. However, drug-induced side effects and poor patient survival hampers the effectiveness of sorafenib chemotherapy (Shikdar, Barry et al.; Cheng, Kang et al. 2009). Our discovery of Utt-B, a saponin derived from the black nightshade (*Solanum nigrum* Linn), as a candidate drug against HCC, have attracted global recognition by obtaining multi-national patents (USA (US2019/0160088A1), Canada (3,026,426.), Japan (JP2019520425) and South Korea (KR1020190008323)), commercial technology transfer (Q Biomed) and above all, gaining the ‘orphan drug’ status by the US FDA, demanding the necessity for further studies to advance this drug to clinics as a treatment option for HCC (R Nath, Swetha et al.; Nath, Gorantla et al. 2016). The current study intends to compare the *in vitro* and *in vivo* chemotherapeutic efficacies of Utt-B with that of sorafenib. Our data highlights the therapeutic supremacy of Utt-B over sorafenib in a murine model of human HCC.

## MATERIALS AND METHODS

### Chemicals

Collection, isolation, and purification of Utt-B were done as previously reported (Nath, Gorantla et al. 2016). Important cell culture reagents such as Dulbecco’s Modified Eagle Medium (DMEM) (GIBCO, 12800-017), Minimum Essential Medium (MEM) (61100061), Foetal Bovine Serum (10270-106), and streptomycin sulfate (GIBCO, 11860-038) were obtained from Invitrogen Corporation (Grand Island, USA). Poly Excel HRP/DAB detection system universal kit (PathnSitu Biotechnologies Pvt. Ltd, India, OSH001) was used for immunohistochemistry experiments. MTT reagent was purchased from TCI Chemicals (India) Pvt. Ltd (D0801). DAPI (D9542), Propidium Iodide (P 4170), RNase A (10109142001), antibodies against β-actin (12620S), GAPDH (8884S), Caspase 9 (9508S), Caspase 7 (12827S) PARP (9532S), and cleaved PARP (5625S) were obtained from Cell Signaling Technologies (Beverly, MA, USA). Annexin V apoptosis detection kit (sc4252AK) and the antibodies against PCNA (sc25280) and Ki67 (sc23900) were purchased from Santa Cruz Biotechnology (Santa Cruz, CA, USA). DeadEnd™ Colorimetric TUNEL System was procured from Promega (G7132). All other chemicals were purchased from Sigma Chemicals (St. Louis, MO, USA) unless otherwise mentioned.

### Cell culture

The liver cancer cell line, HepG2 was purchased from ATCC, Huh-7 from NIH, Hep3B, SK-HEP-1, and immortalized normal Chang Liver cells from NCCS, Pune. MTT assay was performed in cancer lines as previously described (Bava, Puliappadamba et al. 2005). Chang Liver cells were cultured in Minimum Essential Medium (MEM) supplemented with 10% Foetal Bovine serum. All other cell lines HepG2, Huh-7, Hep3B and SK-HEP-1 were cultured in Dulbecco’s Modified Eagle Medium (DMEM) supplemented with 10% Foetal Bovine serum.

### *In vitro* assays

Immunoblotting, immunofluorescence, clonogenic and wound healing assays were performed as previously described (R Nath, Swetha et al.; Vinod, Nair et al. 2015). Immunoblotting analysis was performed as described previously (Saikia, Retnakumari et al. 2018). Immunofluorescence microscopy for the detection of nuclear condensation and apoptosis, using DAPI and Annexin V respectively were performed as previously described (Shankar G, Alex et al. 2020).

### Animals

*In vivo* experiments were conducted according to the RGCB animal ethical committee approval (IAEC/634/RUBY/2017, IAEC/810/RUBY/2020). Mice were housed in a 12h light/dark cycle with access to food and water. Tumor xenografts were generated by the subcutaneous injection of 5×10^6^ HepG2 cells in matrigel into the right lower flank of 6-week-old NOD-SCID (NOD.CB17-Prkdcscid/J) male mice. Two weeks post-injection, the mice were separated into 3 groups (n=6). Group I received the vehicle, Group II and Group III received an intraperitoneal injection of Utt-B in PBS (10 mg/kg body weight) and sorafenib in cremophor vehicle (35 mg/kg body weight) respectively, on alternate days for 1 month, followed by the euthanasia of animals and subsequent procurement of the tissue samples for further analyses. Histology, immunohistochemistry, and TUNEL assay were performed in the tumor samples as previously described (Sreekanth, Bava et al. 2011). The *in vivo* toxicity analysis (acute and sub-chronic) of sorafenib was conducted in 6-8-week-old Swiss albino mice (male and female, 1:1). The test group was administered with varying concentrations of sorafenib (ranging from 0-140 mg/kg body weight). After the culmination of the study period, the liver function and renal profile of the mice were analysed.

### Flow cytometry

FACS and cell cycle analysis were performed as previously reported using BD FACS Diva software Version 5.0.2 (Vinod, Antony et al. 2013).

### Statistical analysis

Data analysis was performed using GraphPad Prism software Version 8 (GraphPad Software Inc., San Diego, CA, USA). *p* < 0.05 was considered as statistically significant. The error bars represent ± SD and are indicative of three independent experiments.

## RESULTS

We previously demonstrated the potency of Utt-B to drive liver cancer cells to death under *in vitro* conditions (Nath, Gorantla et al. 2016). An evaluation of sorafenib under identical conditions revealed the comparative efficacies of both the drugs in inducing liver cancer cytotoxicity. Amongst the various liver cancer cell lines that were employed for the screening, HepG2 was the most sensitive to both the drugs in a cell viability assay and the comparison of their IC50s (0.5 μM and 5.8 μM respectively) indicated that Utt-B is approximately 10-fold potent than sorafenib (Fig. 1A, Fig. S1A, Fig. 1B). Our previous report revealed that Utt-B is not cytotoxic to normal immortalized hepatocytes, Chang Liver, even at elevated drug doses (Nath, Gorantla et al. 2016). However, treatment using high concentrations of sorafenib resulted in the death of Chang Liver, indicating that Utt-B is safer than sorafenib to normal hepatocytes (Fig. S1B). HepG2-induced colony formation was drastically inhibited by Utt-B than sorafenib, indicating the superior anti-clonogenic potential of Utt-B (Fig. 1C). A comparison of HepG2 cell morphologies post-Utt-B and -sorafenib treatment indicated that both the drugs induce apoptotic mode of cell death in HCC cells and that Utt-B is a stronger apoptosis-inducer than sorafenib (Fig. 1D). A similar observation was made in wound healing assay where Utt-B-treated cells demonstrated a reduced wound closure than the sorafenib-treated group, suggesting that Utt-B possesses a better anti-migratory potential over sorafenib (Fig. 1E). Furthermore, flow cytometry analysis revealed that Utt-B drove a significant number of cells to apoptosis in comparison to sorafenib (Fig. 1F). In addition, Utt-B potentiated more cleavage of procaspases 9, 7, and PARP than sorafenib (Fig. 1G, H, I).

**Figure 1.**
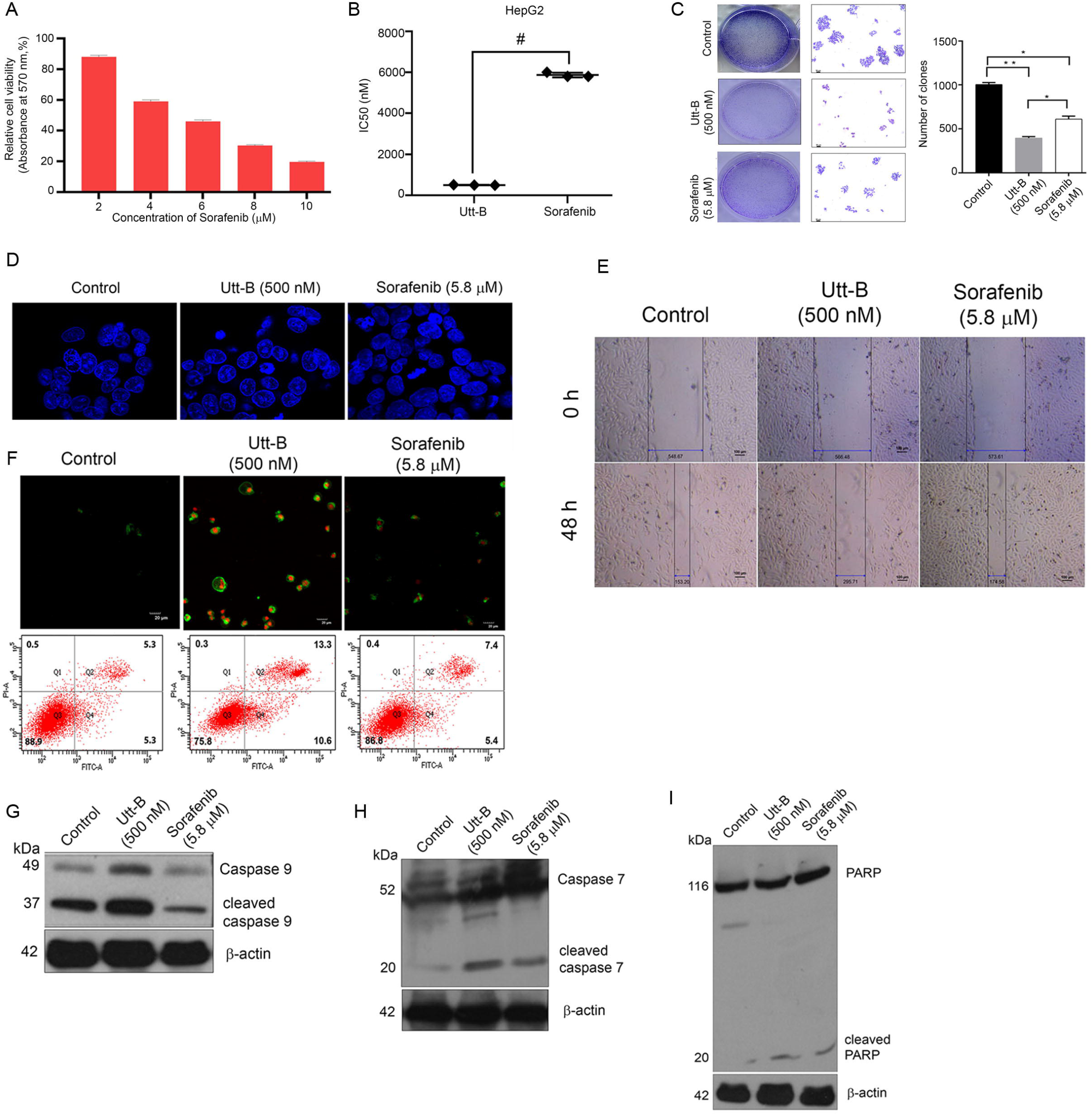
Utt-B induces enhanced apoptosis of HCC in comparison to sorafenib, *in vitro*. **(A)** Assessment of sorafenib cytotoxicity in HepG2 cells, as assessed by MTT Assay. **(B)** A plot between Utt-B *vs* sorafenib treatment, *in vitro*. **(C)** Clonogenic assay reveals an augmented anti-clonogenic potential of Utt-B than that of sorafenib. **(D)** DAPI staining indicating that Utt-B enhances the nuclear condensation of HepG2 cells, than sorafenib. **(E)** Wound healing assay showing the augmented anti-migratory potential of Utt-B than that of sorafenib. **(F)** Annexin-PI flow cytometric analysis shows an increase in apoptosis of HCC cells upon Utt-B treatment. **(G-I)** Immunoblot analysis demonstrates an enhancement in cleavage of caspase 9, 7, and PARP in HepG2 cells treated with Utt-B, in comparison to sorafenib.

A critical xenograft study involving Utt-B and sorafenib revealed their efficacies *in vivo*. The schematic of human HCC-induced tumour development and drug treatment regimen in the NOD-SCID murine model has been provided in Fig. S2 and detailed in the methodology. Following the drug treatment, the tumours were procured for further analysis. Firstly, the tumour volume in the Utt-B- and sorafenib-treated mice was drastically reduced in comparison to the control. However, a striking variation in the reduction of tumour sizes was noticed upon comparing Utt-B- and sorafenib-treated groups, suggesting that Utt-B displays an upper hand in the destruction of HCC cells (Fig. 2A and 2B). Utt-B treatment accentuated caspase 7 cleavage, a marker of the apoptotic programme (Fig. 2C). Histopathological analysis of the excised tumour samples from the various groups authenticated the above observations (Fig. 2D). Furthermore, TUNEL staining revealed the presence of a higher number of apoptotic cells in the Utt-B-treated in comparison to the sorafenib-treated mice (Fig. 2E). The immunohistochemical analysis in the tumour tissues for the expression status of proliferation markers, PCNA and Ki67, and the apoptosis marker, cleaved PARP revealed the augmented potency of Utt-B over sorafenib (Fig. 2F).

**Figure 2.**
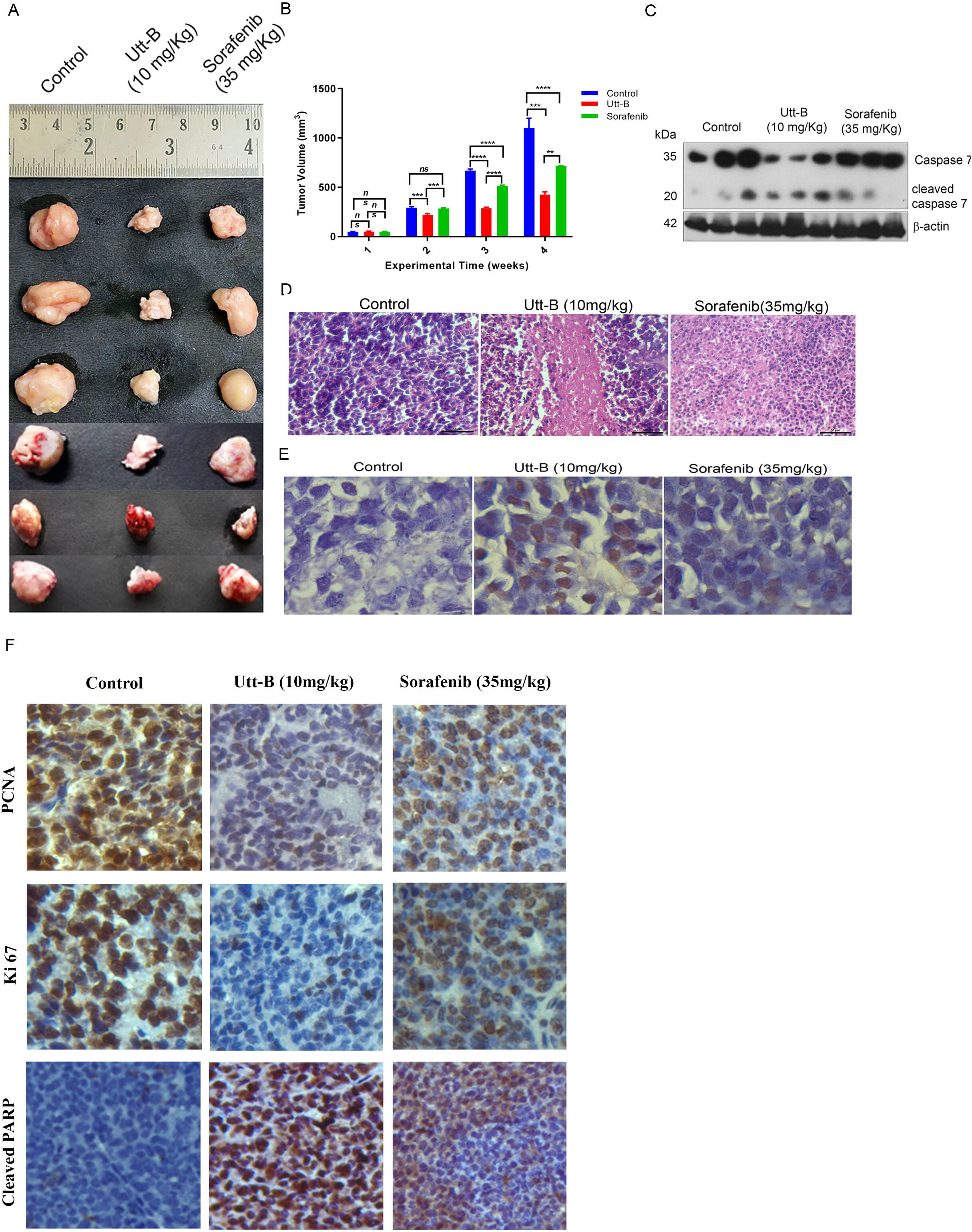
Augmented anti-HCC efficacy of Utt-B over sorafenib, in a NOD-SCID model of human HCC. **(A)** An image of xenograft-excised tumours post-drug treatment regimen. **(B)** A graphical representation of tumour volumes of indicated treatment groups. **(C)** Immunoblot analysis in tumour samples demonstrates an enhanced cleavage of caspase 7 upon Utt-B treatment, in comparison to sorafenib-treated. **(D)** Histopathological staining of xenograft-derived tumours from treatment groups. **(E)** Increase in TUNEL-positive cells following Utt-B treatment, confirms the augmentation of apoptosis. **(F)** Immunohistochemical analysis of nuclear proliferation markers PCNA, Ki67 and the apoptosis marker, cleaved PARP in tumour sections.

We previously reported a detailed toxicological evaluation of Utt-B *in vivo*, demonstrating its pharmacological safety in Swiss albino mice (Nath, Gorantla et al. 2016). Though a well-studied chemotherapeutic drug, we conducted a toxicity evaluation of sorafenib under identical conditions, since some mice in the sorafenib-treated group of the xenograft study exhibited toxicity symptoms such as paralysis of legs and faecal blood stains towards the end of the treatment period. A schematic of toxicity analysis upon sorafenib regimen in Swiss albino mice is depicted in Fig. S1C. However, the analysis of acute and sub-chronic liver toxicity studies revealed that the indicated dosages of sorafenib neither inflicted any behavioural changes nor any shifts in the liver function or renal profile, in comparison to the untreated control mice, attesting the biological safety of sorafenib in normal Swiss albino mice, at the IC50 concentration (Fig. S1D-G). Hence, the previously observed toxicological symptoms in the xenograft study may be attributed to the immune deficiency of NOD-SCID mice. Similar observations have been reported in CB17/16 SCID mice (Kuczynski, Lee et al. 2015). We also noted severe breathing problems, signs of hypertrophy, and regenerative changes in liver hepatocytes in Swiss albino mice treated with four times the IC50 dose of sorafenib (Fig. S1H). On the contrary, only reversible microvesicular fatty changes associated with chemotherapy were observed in liver tissues of the mice treated with even up to five times the IC50 dose of Utt-B in our previous study(Nath, Gorantla et al. 2016). Together, our data demonstrate the superior therapeutic efficacy which Utt-B holds over sorafenib, in the attenuation of HCC.

## DISCUSSION

The present study highlights Utt-B as a better therapeutic agent in comparison with sorafenib, the standard drug used for treating HCC. Sorafenib is widely studied and is known to induce side effects that often result in poor patient survival or recurrence of HCC (Hussaarts, van Doorn et al. 2021; Nasser, El-Naggar et al. 2021). Our evaluation of the sorafenib-associated toxicological parameters in Swiss albino mice revealed that the drug is safe at a dose corresponding to its *in vitro* IC50 concentration. However, higher dosages of sorafenib resulted in severe toxicity in mice as per the acute and sub-chronic toxicity studies. On the contrary, we have already shown that Utt-B is safe to mice even at elevated drug concentrations, authenticating the pharmacological safety of Utt-B over sorafenib. Since Utt-B has been granted the status of an ‘orphan drug’ against HCC by the US FDA, the current study was critical to ensure a comparative screening of this drug with a prominent FDA-approved HCC drug, sorafenib. The human liver cancer cell line, HepG2 was used as a tool for the *in vivo* HCC model, due to its enhanced sensitivity towards both drugs, as per our previous observations (R Nath, Swetha et al.; Nath, Gorantla et al. 2016). Our data indicate that Utt-B possesses all credibility to function as a better anti-HCC drug than sorafenib. Our very recent study has unravelled, with mechanism-based evidence, better modes to further improve the chemotherapeutic potential of Utt-B against HCC (R Nath, Swetha et al.). A systematic clinical evaluation of Utt-B in HCC patients is the next key step to advance this drug from bench to bedside.

## Acknowledgements

We acknowledge DST-SERB for funding. We also acknowledge the immense help provided by the RGCB animal house and instrumentation facilities for the successful completion of the experiments.

## ADDITIONAL INFORMATION

### Ethics approval and consent to participate

Studies involving experiments with animals were conducted in accordance with institution guidelines, under the approval from Institutional Animal Ethics Committee, Rajiv Gandhi Centre for Biotechnology. (CPCSEA Number:326/GO/ReBiBt/S/2001/CPCSEA). The cell lines used in this study were obtained from ATCC.

