## Supplementary Information for "Augmented efficacy of uttroside B over sorafenib in a murine model of human hepatocellular carcinoma"

### Fig. S1.

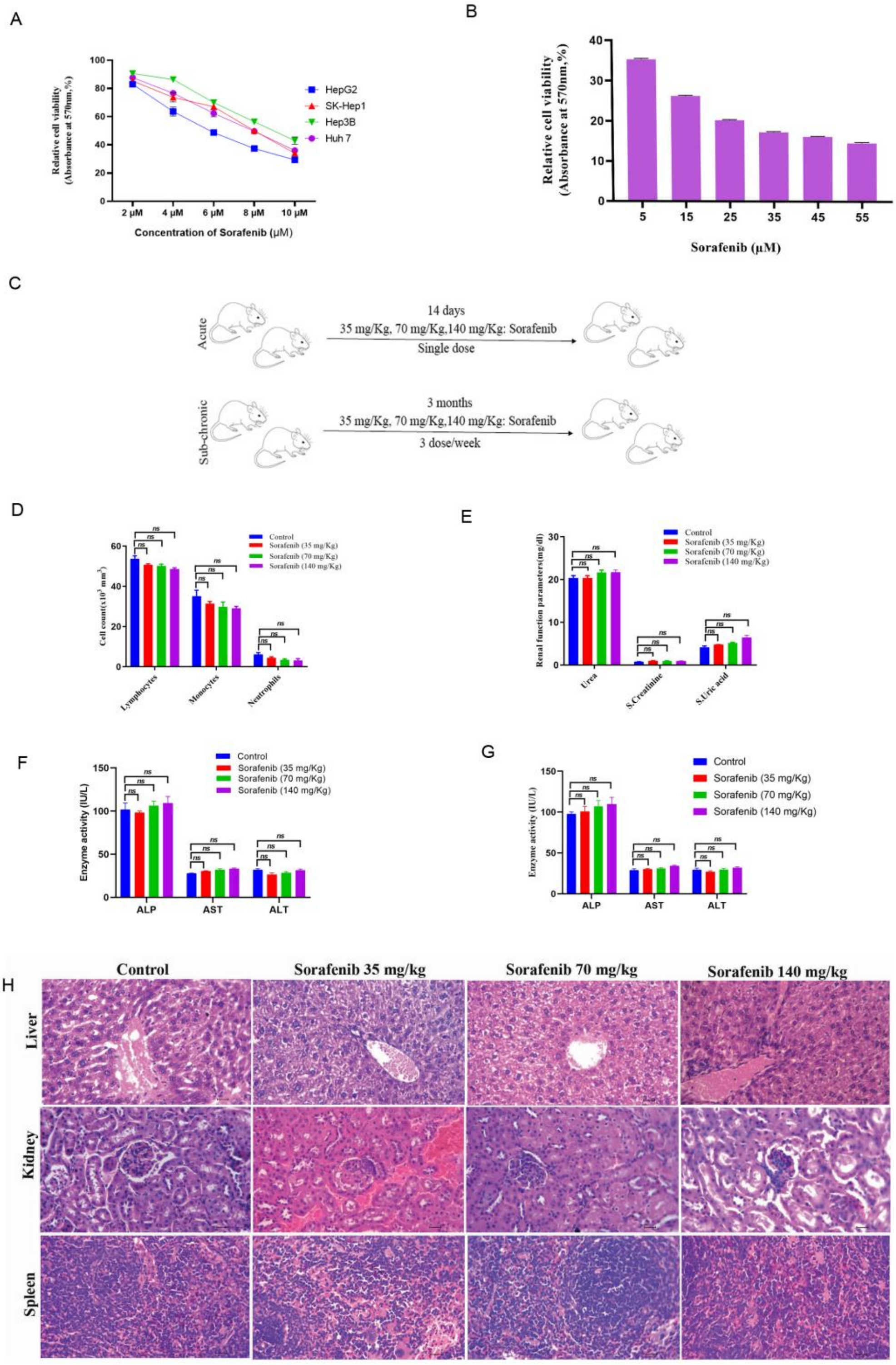

**Fig. S2.**

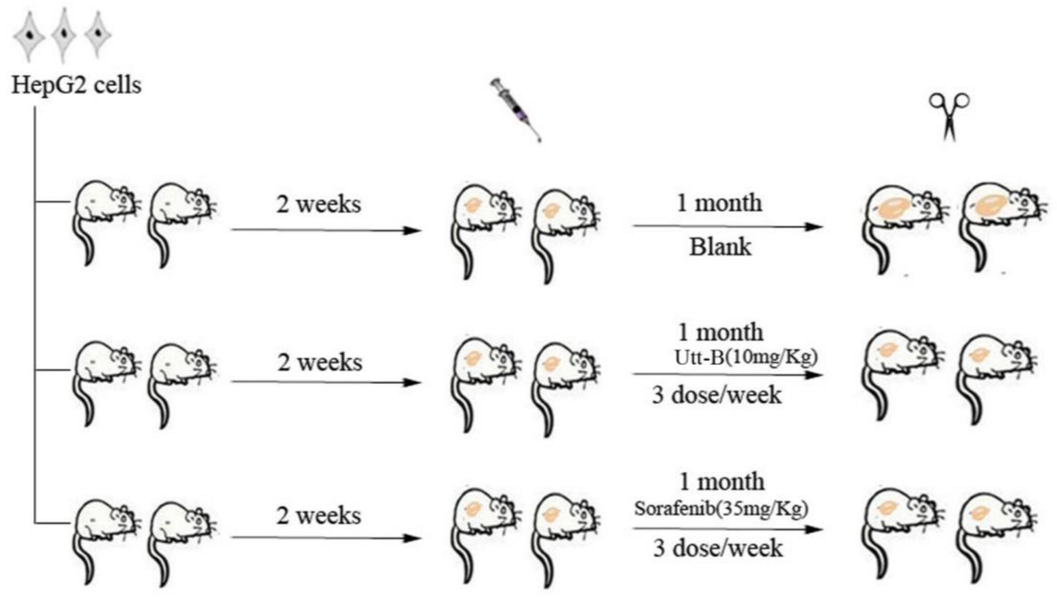


**Fig. S1. Toxicity analysis of sorafenib in Swiss albino mice. (A)** MTT assay of sorafenib in the indicated liver cancer cell lines. **(B)** Cytotoxicity analysis of sorafenib in normal hepatocytes, Chang Liver. **(C)** A scheme of toxicity analysis of sorafenib, in Swiss albino mice. **(D-G)** Acute and sub-chronic toxicity analysis of sorafenib-treated mice groups. **(H)** Histopathological analysis of liver, kidney and spleen tissues of mice post-sorafenib regimen.

**Fig. S2. A scheme depicting the induction of human HCC tumours in NOD-SCID mice using HepG2 cells.**
